# Biomimetic sponges improve muscle structure and function following volumetric muscle loss

**DOI:** 10.1101/2020.05.20.106823

**Authors:** Gabriel Haas, Andrew Dunn, Josh Madsen, Peter Genovese, Andrew Lin, Hannah Chauvin, Jeffrey Au, Allison Paoli, Koyal Garg

## Abstract

Skeletal muscle is inept in regenerating after traumatic injuries such as volumetric muscle loss (VML) due to significant loss of basal lamina and the resident satellite cells. Currently, there are no approved therapies for the treatment of muscle tissue following trauma. In this study, biomimetic sponges composed of gelatin, collagen, laminin-111, and FK-506 were used for the treatment of VML in a rodent model. We observed that biomimetic sponge treatment improved muscle structure and function while modulating inflammation and limiting the extent of fibrotic tissue deposition. Specifically, sponge treatment increased the total number of myofibers, type 2B fiber cross-sectional area, myosin: collagen ratio, myofibers with central nuclei, and peak isometric torque compared to untreated VML injured muscles. As an acellular scaffold, biomimetic sponges provide a promising “off-the-shelf” clinical therapy for VML.

## Introduction

Skeletal muscle is endowed with a remarkable capacity for regeneration, but “volumetric muscle loss” (VML) presents a unique challenge due to the unrecoverable loss of basal lamina and resident stem cell population that results in chronic functional impairment and disability [1–4]. Autologous tissue grafts from an uninjured site are currently used to treat VML injuries in the clinic [5]. However, this complicated surgical approach can cause donor site morbidity, infection, and has been reported to result in graft failure in 4-7% of cases [6, 7]. Physical therapy is typically recommended to VML patients to strengthen the remaining muscle mass, but it is unable to facilitate appreciable muscle regeneration within the site of injury [8]. Therefore, there is a clinical need to develop bioengineered therapies for muscle regeneration and reconstruction following VML.

The extracellular matrix (ECM) forms an ideal microenvironment for cell survival and activity. The ECM not only provides a framework for structural and mechanical support but also sequesters cytokines and growth factors to orchestrate cell migration, proliferation, and differentiation [9, 10]. As a result, acellular ECM scaffolds have been extensively used for the treatment of skeletal muscle injuries. However, there is mounting evidence to suggest that decellularized ECM scaffolds do not support muscle regeneration and remodel into a dense collagenous scar [11–14]. This inability to promote muscle regeneration is primarily attributed to insufficient satellite cell proliferation and activity in the defect region [1, 15, 16]. Furthermore, these scaffolds are unable to modulate the overwhelming immune response that dysregulates the muscle regeneration process [14]. We have previously demonstrated that heightened and prolonged immune response to musculoskeletal trauma impairs regeneration [17]. As early as three days post-VML, both innate and adaptive immune cells can be observed in the VML defect, and their persistent presence impairs muscle regeneration by extending the inflammatory phase and delaying the repair and remodeling phase.

The fungal macrolide FK-506 (Tacrolimus) is an FDA approved immunosuppressant that disrupts signaling events mediated by calcineurin in T lymphocytes [18]. By inhibiting calcineurin, FK-506 prevents dephosphorylation of nuclear factor of activated T cells (NFAT), which reduces the production of IL-2, a cytokine that promotes autocrine T-cell development and proliferation. Besides T-cells, FK-506 can also modulate the pro-inflammatory cytokine production by dendritic cells and macrophages [19, 20]. Since FK-506 selectively reduces pro-inflammatory cytokine production, it can potentially promote T-helper 2 (Th2) versus T helper 1 (Th1) responses [21, 22]. In a recent study, intraperitoneal (i.p.) injection of FK-506 (1 mg/kg) at the time of injury in rats with composite muscle-bone trauma attenuated the immune response in the VML injured skeletal muscle. At day three post-injury and FK-506 injection, total T-cell numbers in VML injured muscle reduced by 67%, helper T-cells specifically by 51%, and macrophages by 22% [23]. However, the administration of FK-506 alone did not result in functional muscle regeneration. In a subsequent study, when FK-506 was injected i.p. and VML defect was treated with autologous minced muscle grafts in a porcine model, muscle regeneration and function was enhanced [24]. Taken together, these studies suggest that a combination of regenerative and immunomodulatory therapy is essential for functional muscle regeneration.

In a previous study, we demonstrated that a unique blend of gelatin, collagen, and laminin-111 in a porous sponge-like scaffold supports the infiltration of satellite cells, promotes myogenic protein expression, and myofiber regeneration in a murine model of VML [25]. In this study, we incorporated FK-506 in the biomimetic sponges to enhance their immunomodulatory properties. We hypothesize that biomimetic sponges will support functional recovery in a rat model of VML by stimulating regeneration and limiting the extent of inflammation and fibrosis.

## Materials and Methods

### Preparation of collagen/gelatin/laminin/FK506 sponge

A 3 wt % porcine skin gelatin (Sigma-Aldrich) solution was prepared in DI water and heated to 60°C. After the gelatin had completely dissolved, the solution was allowed to cool to 50°C. EDC (20 mM) was added to the solution which was combined with rat tail collagen I (Gibco, 3 mg/mL) at a 70:30 gelatin : collagen ratio in a tube. Laminin (LM)-111 (Trevigen) and FK506 (Abcam) were added to the solution at final concentrations of 50 μg/mL and 25 μM, respectively. The final solution was then aliquoted into a 48-well plate at 700 μL/well. The well plate was placed in a 100% methanol bath, allowing the sponges to gel at 4°C for 30 min, followed by overnight freezing at −8°C. The well plate was then removed from the bath and moved to a −80°C freezer for 48 hours and subsequently lyophilized for at least 12 hours. The day before surgery, the sponges were disinfected via a 5-minute incubation in 200-proof pure ethanol, which was followed by two 5-minute rinses, and an overnight rinse in sterile 1x phosphate-buffered saline (Gibco).

### Implantation of sponges into a VML model

This work was conducted in compliance with the Animal Welfare Act, the implementing Animal Welfare Regulations, and in accordance with the principles of the Guide for the Care and Use of Laboratory Animals. All animal procedures were approved by the Saint Louis University's Institutional Animal Care and Use Committee.

Male Lewis rats (10-12 weeks old) were purchased from Charles Laboratory and housed in a vivarium accredited by the Association for Assessment and Accreditation of Laboratory Animal Care International and provided with food and water ad libitum. The animals were weighed prior to surgery and anesthetized using 2.5% isoflurane. The surgical site was aseptically prepared, and sustained release buprenorphine (1 mg/kg) was injected in the nape of the neck prior to the procedure. A lateral incision was made through the skin to reveal the tibialis anterior (TA), and the skin was separated from the musculature by blunt dissection. A metal plate was inserted underneath the TA muscle, and a 6-mm punch biopsy was performed to remove approximately ~20% of the muscle mass. The biopsy was removed and weighed for consistency. A subset of injured animals received biomimetic sponges while the other subset was left untreated. A 6-mm sponge disk was used to treat the TA. Bleeding was controlled with light pressure, and the skin incision was closed with simple interrupted skin staples. The animals were allowed to recover for 7, 14, or 28 days and euthanized via exsanguination followed by cervical dislocation. TA muscles were weighed upon collection (n = 9 – 17 muscles, and 5 – 9 animals per group) and processed for histological and biomolecular analyses. The surgery was performed bilaterally, keeping the treatment subsets consistent between both legs.

### Histology

TA muscles were cut in half at the defect site, and the upper portion was frozen in 2-methylbutane (Fisher Scientific) super-cooled in liquid nitrogen for 10 seconds. The muscles were mounted on stubs using OCT, and transverse cross-sections (15 μm) were cryosectioned from the area where the original surgical defect was made. Cross-sections were stained with hematoxylin and eosin (H&E), collagen 1 (1:100; ab34710; Abcam, Cambridge MA), sarcomeric myosin (1:50; MF20; Developmental Studies Hybridoma Bank), laminin (1:100; ab11575; Abcam; Cambridge MA), CD68 (1:50; MCA341R; AbD Serotec, Raleigh, NC), nuclei (DAPI; 1:100; Invitrogen), CD3 (1:100; ab5690; Abcam, Cambridge MA), CD31 (1:100; R&D systems), α-bungarotoxin (1:100; Invitrogen). Appropriate fluorochrome-conjugated secondary antibodies (1:100; Invitrogen) were used as described previously [1, 3, 5]. Images were captured at 10× and 20× magnification using a Zeiss Axiocam microscope. Slides were scanned to obtain composite images of the entire muscle section using Olympus BX614S (Saint Louis University) and NanoZoomer 2.0 HT (Washington University in Saint Louis).

Full-size muscle sections stained with H&E were used to quantify the number of myofibers with centrally located nuclei at day 28 post-injury (n = 4 muscles, 2 – 3 animals). Full-size muscle sections stained with myosin heavy chain (MHC) and collagen (COL) were used to quantify the MHC:COL ratio by % area (n = 4 – 5 muscles, 2 – 4 animals). Images were opened in ImageJ (or Fiji) and the remaining defect area and remaining muscle tissue were divided into two separate images of the same size. The two images (i.e., one containing the defect area and the other containing the remaining healthy tissue), were separately analyzed by splitting the color channels, thresholding the MHC and COL to most accurately represent the stained area, and measuring the percentage of area positively stained by MHC and collagen. The % area of MHC and collagen from the two images were added up, and the MHC: COL ratio was determined. The muscle sections were split into remaining muscle tissue and defect area in order to measure the collagen deposition accurately. The collagen tissue deposited in the defect showed different sensitivity to thresholding than the collagen in the remaining healthy tissue. When thresholding without splitting, low thresholding results in the defect area not being picked up, whereas high thresholding results in collagen filling the muscle fibers. Thresholding the defect region and the remaining healthy tissue separately allowed for accurate detection and measurement of collagen (Supplemental Figure 1).

Muscle cross-sections were stained using antibodies from Developmental Studies Hybridoma Bank (Iowa City, IA, USA) for fiber types 1 (1:20; BA.D5), 2A (1:50; SC.71), and 2B (1:20; BF.F3), as described previously [26]. A laminin counterstain (1:100; ab11575) served as the fiber outline. Unstained fibers were identified as type 2X. A custom-designed image analysis MATLAB program was used for the quantification of myofiber cross-sectional area (CSA; n = 4 – 5 muscles, 3 – 4 animals) and fiber type distribution (n = 3 – 5 muscles, 3 – 4 animals). Briefly, the algorithm first thresholds the laminin channel, followed by nonlinear morphological transformations to delineate fiber boundaries and reduce noise. Area, circularity, and concavity filters are applied to identify myofibers. To avoid spatial variances in brightness, a color histogram is computed for each fiber, and areas corresponding to only a single channel are then compared to determine the primary fiber-type. If no color is dominant, the fiber is marked as 2X.

### Flow Cytometry

Cells were isolated from the entire TA muscle with the muscle defect (n = 4 – 6 muscles, 2 – 4 animals) by enzymatic digestion as previously described [17, 23]. Briefly, the TA was surgically isolated, and the mass was determined. Tissue was incubated with collagenase type II and dispase for 90 min at 37 °C. Cells were further released by gentle mechanical disruption and filtered through a 70 μm cell strainer and subsequently through a 40 μm cell strainer. Erythrocytes were lysed with ammonium-chloride-potassium lysing buffer, and cells were washed and resuspended in Roswell Park Memorial Institute (RPMI) medium. Viable cells were quantified using trypan blue exclusion and a hemocytometer. After quantification, cells were resuspended at 106 cells/mL in PBS containing 0.5% FBS and 0.1% sodium azide. From the resuspension, the cells were incubated with anti-CD32 antibody (BD Biosciences 550271) to block Fc receptors and labeled with either cocktail monoclonal antibodies to identify macrophages or T lymphocytes. The macrophage cell cocktail included anti-CD11b (BD Biosciences 562102, clone WT.5), anti-CD86 (BD Biosciences 743211, clone 24F), and anti-CD163 (Bio-Rad MCA342F, clone ED2). The T lymphocyte cell cocktail (BD Biosciences 558493) consisted of anti-CD3 (IF4), anti-CD4 (OX-35), and anti-CD8α (OX-8). Samples were run on BD biosciences LSR II flow cytometer at the flow cytometry research core facility at Saint Louis University.

### Gene Expression

As described previously [5], RNA was isolated from snap-frozen cross-sections of TA muscle (n = 4 – 5 muscles, 2 – 4 animals) that was comprised of both the defect area and the remaining muscle mass (50 mg). RNA was extracted using Trizol LS reagent (Invitrogen) and purified using RNeasy mini kit (Qiagen). The yield of RNA was quantified using a NanoDrop spectrometer (NanoDrop Technologies) and optical density (OD) 260/280 ratios were determined. RNA (500 ng) was reverse transcribed into cDNA using the Super-Script III first-strand synthesis kit (Invitrogen). Custom designed primers (Sigma-Aldrich) with the sequences presented in Table 1 were used as myogenic and immunogenic markers. All primer sets have been synthesized by Sigma-Aldrich DNA oligos design tool. Aliquots (2 μl) of cDNA were amplified with 200 nM forward/reverse primers, SYBR GreenER (Invitrogen) in triplicate using a Bio-Rad CFX96 thermal cycler system (Bio-Rad). Nontemplate control and no reverse transcriptase controls were run for each reaction. Gene expression was normalized to 18S (housekeeping gene) to determine the ΔCT value. Expression levels for each mRNA transcript were determined by the 2^−ΔΔCT^ method by normalizing each group to uninjured cage control muscles.

**Table 1.**
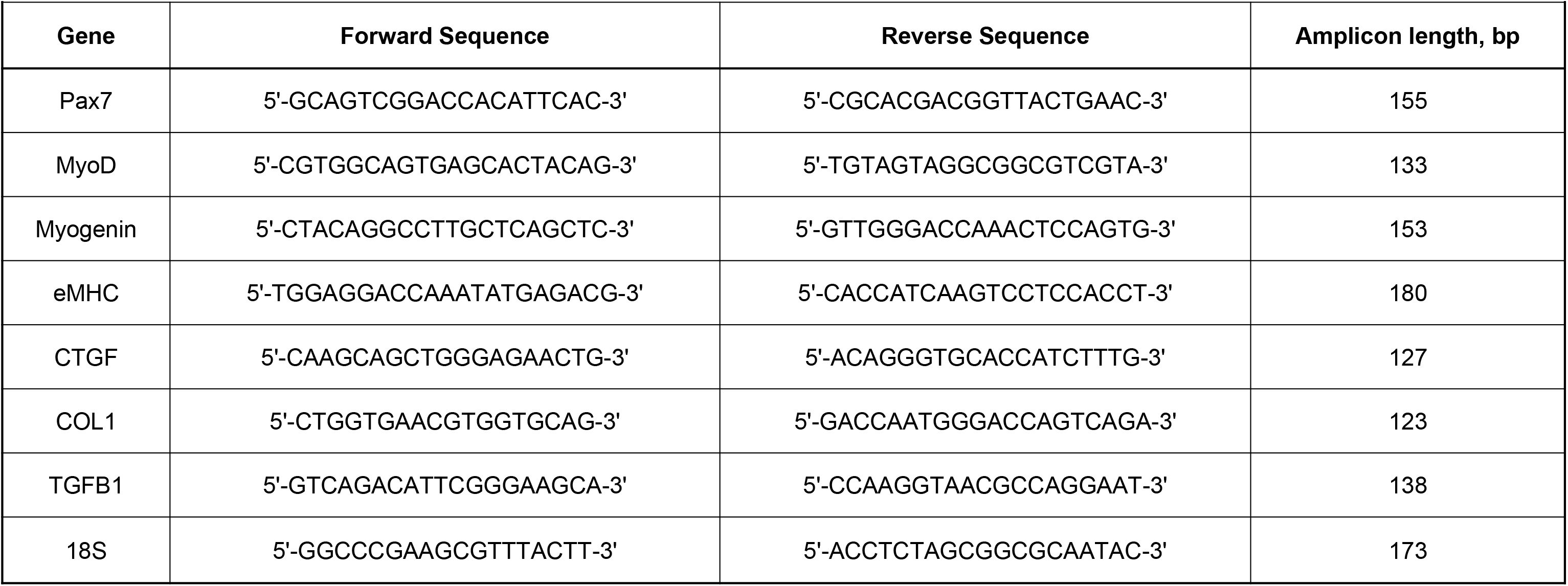
Primers for qrtPCR

Additionally, gene expression of immunogenic markers was measured using a PCR immunoarray, as described previously [17]. RNA isolated as described above and was converted into complementary DNA (cDNA) using RT^2^ First Strand Kit (Qiagen). An RT^2^ Profiler PCR array (PARN052Z, SABiosciences) was employed to examine rat innate and adaptive immune responses. A CFX96 Real-Time System (Bio Rad) was used to determine gene expression. Data analysis was carried out either with the online RT^2^ Profiler PCR Array Data Analysis version 3.5 or expression levels for each mRNA transcript were determined by the 2^−ΔCT^ method. Gene expression in VML-injured muscle tissue was relative to expression in uninjured cage control muscle tissue and normalized to reference gene Rplp1 (n=4 per group).

### Muscle Function Assessment

In vivo functional testing of the anterior crural muscles (n = 7 – 10 muscles, 4 – 6 animals) was performed at 28 days post-injury using the methodology previously described [23]. Briefly, in vivo physiological properties were measured in anesthetized rats (isoflurane 1.5 – 2.0%) using a dual-mode muscle lever system (Aurora Scientific, Inc., Mod. 305b). The skin was shaved, and an incision was made at the postero-lateral aspect of the ankle. The distal tendon of the gastrocnemius-soleus complex muscles was isolated and severed to prevent plantarflexion. Subcutaneous needle electrodes were inserted in the posterior compartment of the lower limb on each side of the common peroneal nerve. Optimal current (30 – 40 mA) was set with a series of twitches. Isometric tetanic contractions were elicited at 150 Hz (0.1 ms pulse width, 400 ms train) with the ankle at a right angle.

### Statistical analysis

Data are presented as a mean ± standard error of the mean. Analysis and graphing of data were performed using GraphPad Prism 8 for Windows. A one-way or two-way analysis of variance was used when appropriate to determine if there was a significant interaction or main effect between variables. The Fisher’s least significant difference post-hoc comparison was performed to identify significance with p ≤ 0.05. The PCR immunoarray data was analyzed using the non-parametric Kruskal-Wallis test followed by Dunn’s post-hoc test to identify significance with p ≤ 0.05.

## Results

### FK506 Release in vitro

The amount of FK506 released from the sponges over 7 days is shown in Figure 1. The majority of the FK-506 loaded on to the sponges was released within the first 3 days, suggesting a burst release. At days 1 and 2, sponges loaded with 25 μM of FK-506 released a significantly higher percentage of the drug compared to sponges loaded with 50 and 100 μM of FK-506. On day 4, almost all of the FK-506 in the sponges had diffused out *in vitro*.

**Figure 1.**
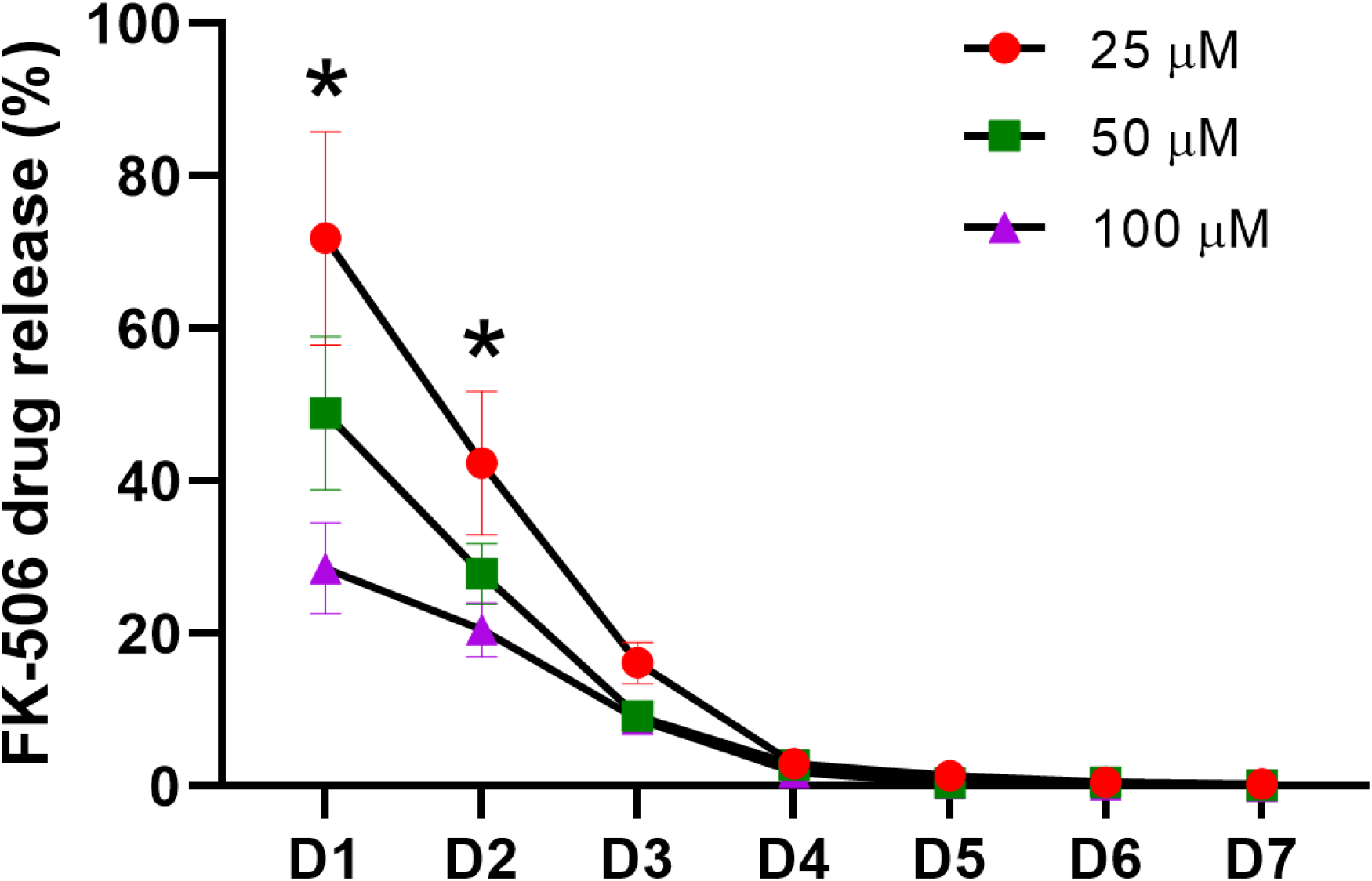
FK506 release from biomimetic sponges was measured *in vitro*. Regardless of initial concentration, most of the drug was released within the first three days. “*” indicates a statistical difference (p < 0.05) in 25 μM sponges versus 50 μM and 100 μM sponges at a particular time-point.

### Myogenic marker expression

Gene expression of myogenic markers is shown in Figure 2. Pax7 and MyoD are markers associated with satellite cell activation and proliferation. Irrespective of treatment, Pax7 shows a transient decrease on day 14 followed by an increase on day 28. MyoD expression shows a linear increase between days 7 and 28 in both treated and untreated samples. Although not significant, myogenin expression is higher with sponge treatment on day 7. Similar to Pax7, it also shows a transient drop at day 14 followed by an increase at the day 28 time-point, but only in the sponge treated group. In the untreated group, myogenin expression remains relatively constant over 28 days. Embryonic myosin heavy chain (eMHC) maintains its expression over 14 days in the sponge treated group. However, it shows a sharp decline in the untreated groups. eMHC expression was higher in the sponge treated group compared to the untreated group at day 14, but statistical significance was not reached.

**Figure 2.**
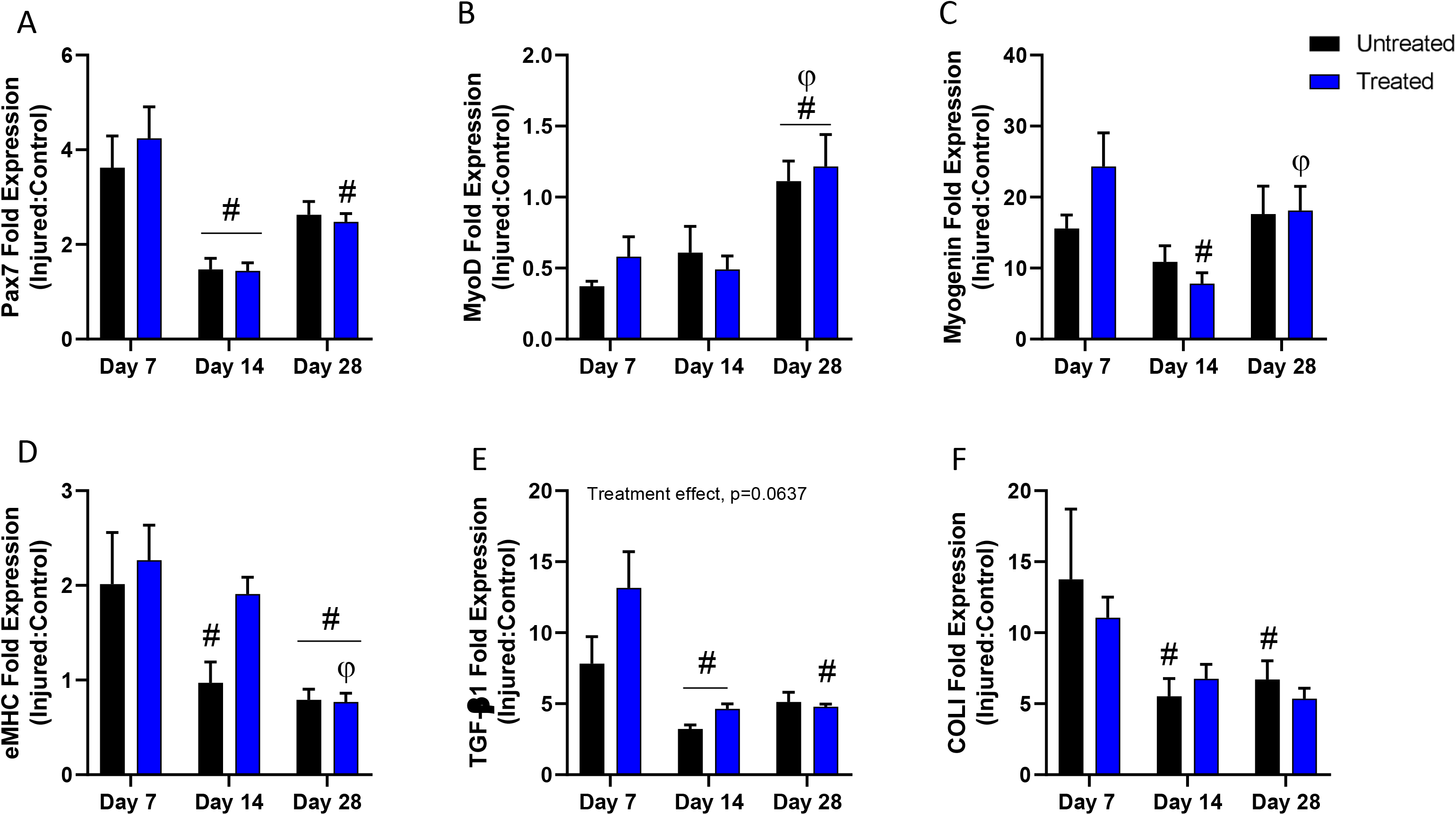
Myogenic gene expression was determined by PCR. The quantification of (A) Pax7, (B) MyoD, (C) Myogenin, (D) eMHC, (F) TGF-β1 and (G) COL1 fold expression is shown for days 7, 14, and 28 post-injury. “#” indicates a statistical difference (p < 0.05) from Day 7 and “φ” indicates a statistical difference (p < 0.05) from Day 14 for a particular treatment group.

A trend towards increased TGF-β1 expression is observed with sponge treatment. Over time, both treated and untreated muscles show a sharp decline in TGF-β1 expression. Collagen 1 gene expression is significantly reduced over time in the untreated muscles, but the expression is maintained in the sponge treated muscles.

### Flow cytometry

Quantitative analysis of mononuclear cell infiltration was performed using flow cytometry at days 7, 14, and 28 post-injury. Cells isolated from the TA muscles were gated for T-lymphocytes (CD3^+^), followed by helper T-lymphocytes (CD4^+^) and cytotoxic T-lymphocytes (CD8^+^). T-lymphocytes could be detected in the VML injured muscles as early as 7 days post-injury (Figure 3). At day 7, no differences in T-lymphocyte quantity were observed between untreated and treated VML injured muscles.

**Figure 3.**
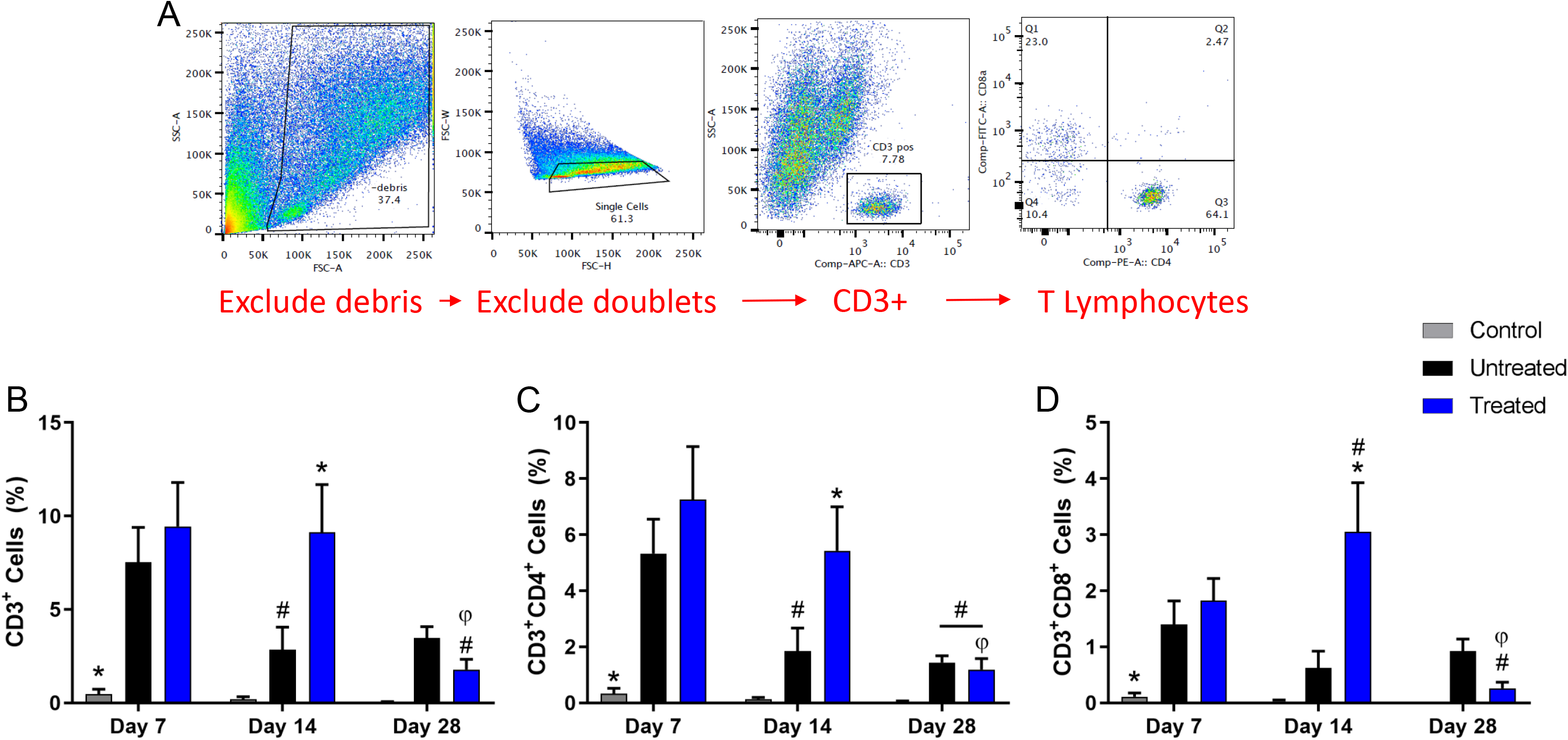
Flow cytometry was used to quantify the infiltrating T-cells. (A) The gating strategy is shown. The percentage of (B) total T-cells (CD3^+^), (C) helper T-cells (CD3^+^ CD4^+^), and (D) cytotoxic T-cells (CD3^+^ CD8^+^) cells were quantified. Injured muscles had higher percentages of T-cells compared to uninjured controls at day 7 post-injury. Treated muscles contained more T-cells than both untreated and uninjured muscles at day 14. By day 28, the T-cell response subsided. “*” indicates a statistical difference (p < 0.05) between different treatment groups at a particular time-point. “#” indicates a statistical difference (p < 0.05) from Day 7 and “φ” indicates a statistical difference (p < 0.05) from Day 14 for a particular treatment group.

On day 14, the untreated muscles showed a significant decrease in both CD3^+^ and CD3^+^CD4^+^ T-cells quantity but not CD3^+^CD8^+^ T-cells. In contrast, the quantity of both CD3^+^ and CD3^+^CD4^+^ T-cells was maintained in the sponge treated muscles over 14 days. The sponge treated muscles showed a significant increase in CD3^+^CD8^+^ T-cell quantity at day 14 compared to day 7. A significantly higher quantity of CD3^+^, CD3^+^CD4^+^, and CD3^+^CD8^+^ T-cells were observed in the sponge treated muscles compared to untreated muscles at day 14. On day 28, no significant differences were observed between treated and untreated muscles. Sponge treated muscles showed a significant decrease in CD3^+^, CD3^+^CD4^+^, and CD3^+^CD8^+^ T-cell quantity compared to untreated muscles. Altogether, these data indicate that while in the untreated muscles the adaptive immune response subsides by day 14, an elevated and persistent T-cell response is observed in sponge treated muscles.

Mononuclear cells were also gated for CD11b^+^ myeloid cells, followed by CD86^+^ pro-inflammatory M1-like cells and CD163^+^ anti-inflammatory M2-like cells (Figure 4). The quantity of CD11b^+^ myeloid cells or that of M1-like (CD11b^+^CD86^+^) and M2-like (CD11b^+^CD163^+^) cells was not significantly different between treated and untreated groups on days 7, 14, and 28. A linear decrease in cell quantity was observed in both treated and untreated samples over 28 days. Overall, these data show mixed recruitment of pro- and anti-inflammatory macrophages in response to VML injury and sponge treatment.

**Figure 4.**
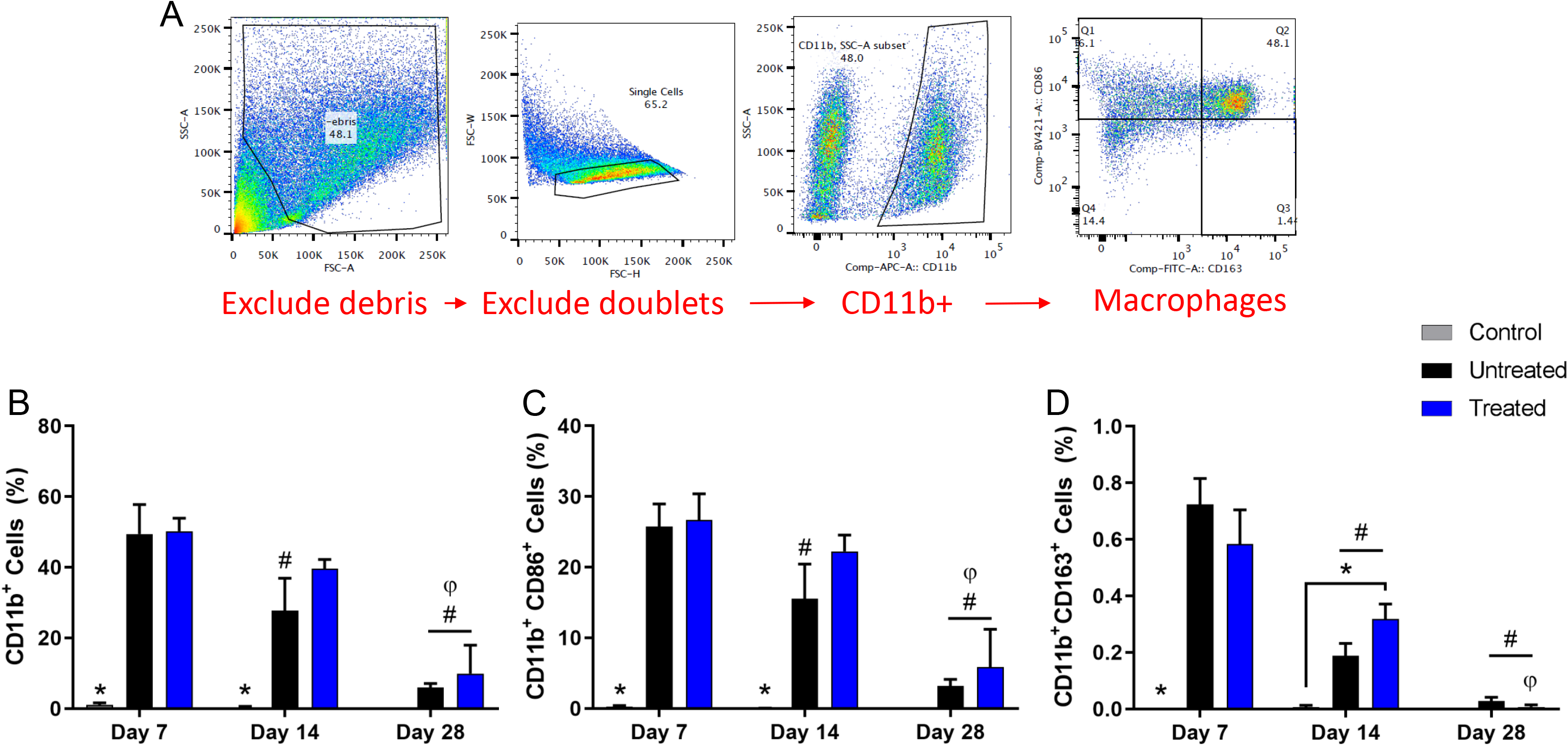
Flow cytometry was used to quantify the infiltrating macrophages. (A) The gating strategy is shown. The percentage of (B) total myeloid cells (CD11b^+^), (C) pro-inflammatory cells (CD11b^+^ CD86^+^), and (D) anti-inflammatory (CD11b^+^ CD163^+^) cells were quantified. At days 7 and 14 post-injury, total myeloid cells and pro-inflammatory cells were elevated in injured muscles. Anti-inflammatory (CD11b^+^CD163^+^) cells were increased in injured muscles at day 7, but only remained elevated in treated muscles at day 14. The myeloid cell response subsided in injured muscles by day 28. “*” indicates a statistical difference (p < 0.05) between different treatment groups at a particular time-point. “#” indicates a statistical difference (p < 0.05) from Day 7 and “φ” indicates a statistical difference (p < 0.05) from Day 14 for a particular treatment group.

### Modulation of the immune response post-VML

A PCR array was used to quantify gene expression of markers associated with innate and adaptive immune responses (Figure 5). VML injury resulted in significant upregulation of toll-like receptors (TLR) −1, 2, 4, and 7 compared to cage controls. Treatment of VML with sponge matrix resulted in significant upregulation of TLR −5, 6, 9, in addition to TLRs-1, 2, 4, and 7 compared to cage controls. In untreated muscles, a trend towards increased gene expression of TLR-6 (p=0.0776) was observed. These results suggest that various TLRs are involved in the innate immune system’s recognition and interaction with the implanted biomaterial [27, 28]. NOD-like receptors (NLR) are a class of pattern recognition receptors that are found in the cytosol. The gene expression of NLRP3 was significantly increased in the sponge treated muscles. The gene expression of NOD2 was trended towards an increase in sponge treated muscles (p=0.0955) but was significantly higher in untreated muscles, compared to cage controls.

**Figure 5.**
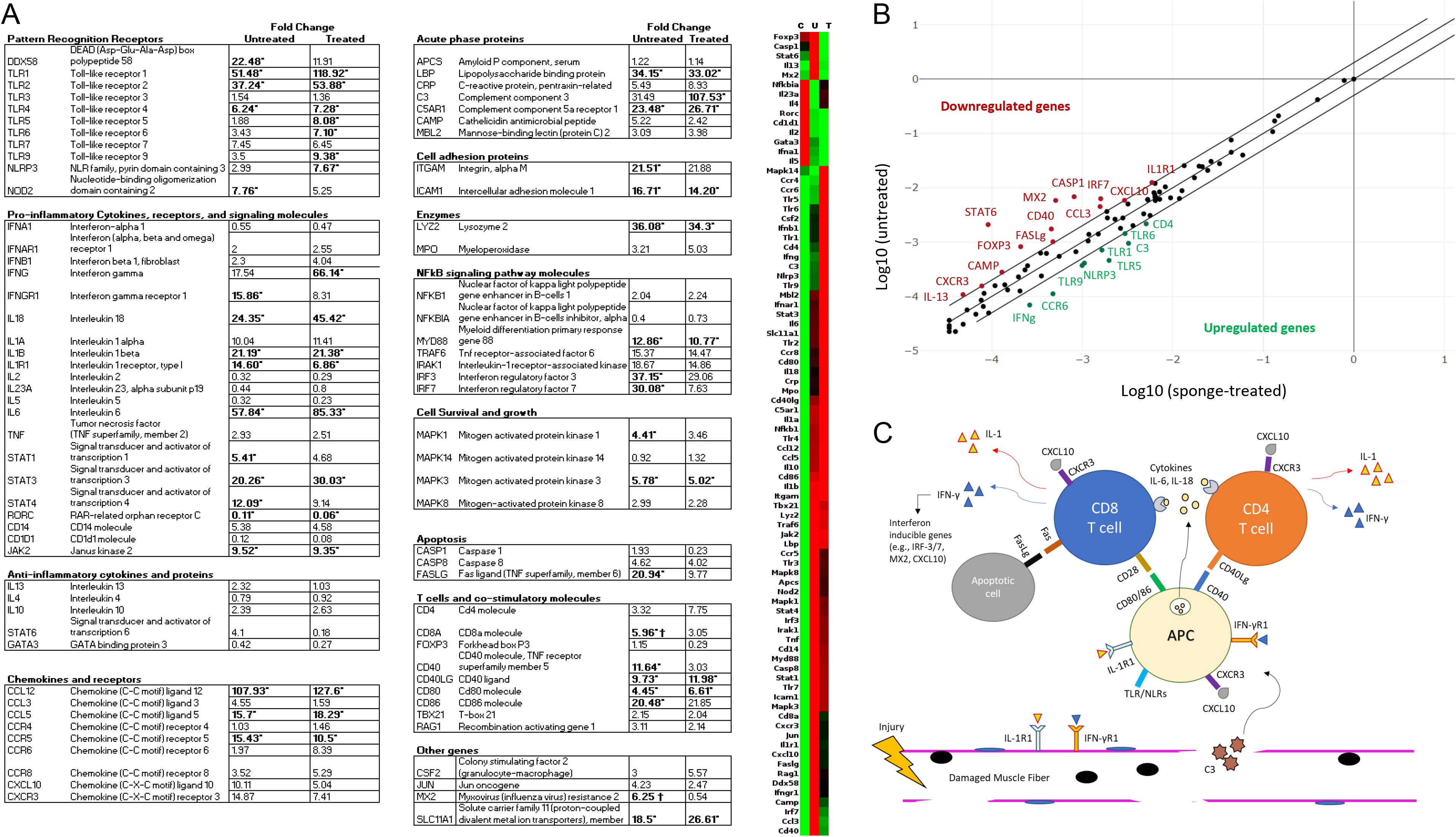
An array of immunogenic markers was quantified via PCR. (A) The table shows specific genes and their fold changes in both untreated and treated muscles at 7 days post-injury. “*” indicates statistical differences (p < 0.05) between injured muscles and controls. “†” indicates statistical differences (p < 0.05) between untreated and treated muscles. (B) Genes downregulated and upregulated with sponge-treatment are shown. (C) A schematic of the implicated immunological events is presented.

Myeloid differentiation factor 88 (MyD88) is an essential adaptor molecule for most TLRs, and it mediates the activation of nuclear factor κB (NF-κB) signaling pathway. Irrespective of treatment, both VML injured muscles showed significantly higher expression of MyD88. Once activated via the NFκB signaling pathway, the innate immune system initiates the inflammatory response secreting chemokines and cytokines. The expression of interferon response factor (IRF)-3 and IRF7 was significantly higher in untreated VML injured muscles compared to cage controls. IRF3 gene expression trended higher in the treated muscles (p=0.0624). As expected, VML injury caused increased gene expression of pro-inflammatory cytokines such as interleukin (IL)-1β, IL-18, and IL-6, irrespective of treatment. Sponge treatment significantly increased the expression of interferon (IFN)-γ. The untreated VML injured muscles showed a significant increase in the expression of cytokine receptor IFN-γR1. While the expression of IL-1R1 was significantly increased in both untreated and treated muscles, its expression was downregulated −2.13 fold in the sponge treated muscles. Sponge treatment resulted in a significantly higher expression of complement component 3 (C3) compared to cage controls.

Several chemokines and their receptors were also significantly increased in response to VML injury such as CCL12, CCL5, and CCR5. Gene expression of both CXCL10 and CXCR3 that are involved in peripheral homing of T-cells was downregulated −2.01-fold with sponge treatment. The chemokine (C-X-C motif) ligand 10 (CXCL10) trended lower in sponge treated muscles (ANOVA p = 0.0575). Cell adhesion proteins such as ITGAM and ICAM1 were increased in both treated and untreated muscles. The gene expression of ITGAM was significantly higher in untreated muscles and trended higher in treated muscles (p=0.0624) compared to cage controls.

The JAK-STAT pathway is involved in the secretion of cytokines. The expression of STAT1 and STAT4 was significantly increased only in untreated muscles but trended higher in the treated muscles (p=0.0955). STAT3 gene expression was significantly increased in both untreated and treated muscles. In response to VML, a significant increase in Jak 2 was observed, irrespective of treatment. Although not statistically different, the gene expression of STAT6 and IL-13 was downregulated −23.04-fold and −2.25-fold, respectively.

Inflammatory factors released by innate immune cells can also recruit T-cells to the site of injury. VML injury resulted in significantly higher gene expression of CD8a, CD40, and CD86 compared to cage controls. Sponge treatment resulted in significantly lower CD8a compared to untreated muscles. Apoptosis associated genes such as Fas ligand (FasL) and caspase 1 were downregulated in sponge treated muscles −2.14 and −8.42-fold, respectively. FasL was significantly increased in untreated muscles compared to cage controls. Both untreated and treated VML injured muscles showed significantly higher expression of CD40 ligand (CD40LG) and CD80. The expression of MX2 was significantly higher in the untreated muscles only. MX2 was downregulated −11.52 fold in the sponge treated muscles compared to untreated muscles.

### Muscle Mass and Function

TA muscle mass was significantly lower in the VML injured muscles irrespective of treatment on days 7, 14, and 28 post-injury compared to the uninjured controls (Figure 6A). A >35% deficit in muscle mass was observed between uninjured controls and VML injured muscles over 28 days. No significant differences were observed between untreated and treated muscles at any time-points.

**Figure 6.**
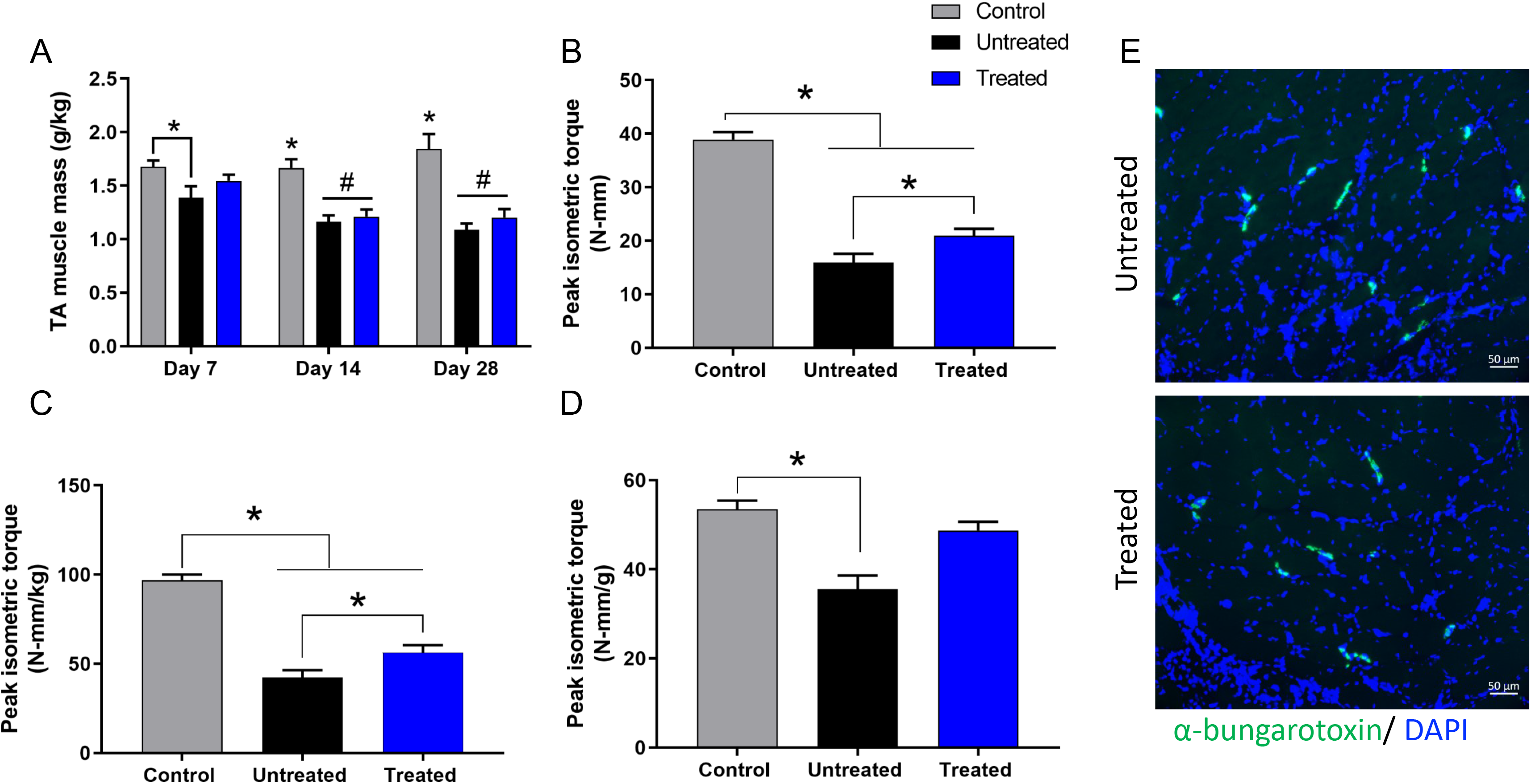
**(A)** Tibialis anterior muscles were weighed upon collection at days 7, 14, and 28 post-injury. (B) Peak isometric torque was measured and (C) normalized to body weight and (D) TA weight. (E) Qualitative analysis of injured muscles showed no differences in the prevalence of post-synaptic terminals. “*” indicates a statistical difference (p < 0.05) between different treatment groups at a particular time-point. “#” indicates a statistical difference (p < 0.05) from Day 7 for a particular treatment group.

The isometric strength of the anterior crural muscles was measured *in vivo* at 28 days post-injury. VML injury resulted in a ~56-59% deficit in peak isometric torque production when analyzing raw force data and that normalized to body mass (Table 2). Sponge treatment significantly improved the torque production in VML injured muscles (Figure 6B-D). Compared to untreated muscles, biomimetic sponges improved muscle strength by ~31-37% when analyzing raw force data and that normalized to body mass or TA muscle mass. Histological analysis of acetylcholine receptor clustering revealed no differences in muscle innervation between untreated and treated muscles (Figure 6E).

**Table 2.** Isometric Torque Production

| Table 2. Isometric Torque Production |  |  |  |  |
| --- | --- | --- | --- | --- |
| Parameters |  | Four weeks post-injury |  |  |
|  |  | Uninjured | Injured |  |
|  |  | Cage Control | Untreated | Treated |
| Unnormalized | Peak Isometric Torque (N-mm) | 38.91 | 15.90 | 20.90 |
|  | Functional Deficit (%) |  | 59.14 | 46.29 |
|  | Functional Recovery (%) |  |  | 31.45 |
| Normalized to Body Weight | Peak Isometric Torque (N-mm/kg) | 96.87 | 42.19 | 56.30 |
|  | Functional Deficit (%) |  | 56.45 | 41.88 |
|  | Functional Recovery (%) |  |  | 33.44 |
| Normalized to TA Weight | Peak Isometric Torque (N-mm/g) | 53.46 | 35.53 | 48.75 |
|  | Functional Deficit (%) |  | 33.54 | 8.81 |
|  | Functional Recovery (%) |  |  | 37.21 |

### Cellular infiltration, myofiber regeneration, and fibrosis

Transverse cross-sections of TA muscles were stained with H&E and are shown in Figure 7. At all three time-points, increased cellular infiltration (DAPI^+^) can be observed in the defect site. At Day 7, the implanted sponges can be identified in the VML defect. The sponges appear to support cellular infiltration and do not cause fibrotic capsule formation. By day 14, the sponges are no longer visible in the defect. On day 28, quantitative analysis revealed that there are significantly more myofibers with centrally located nuclei in treated muscles versus untreated muscles (Figure 7).

**Figure 7.**
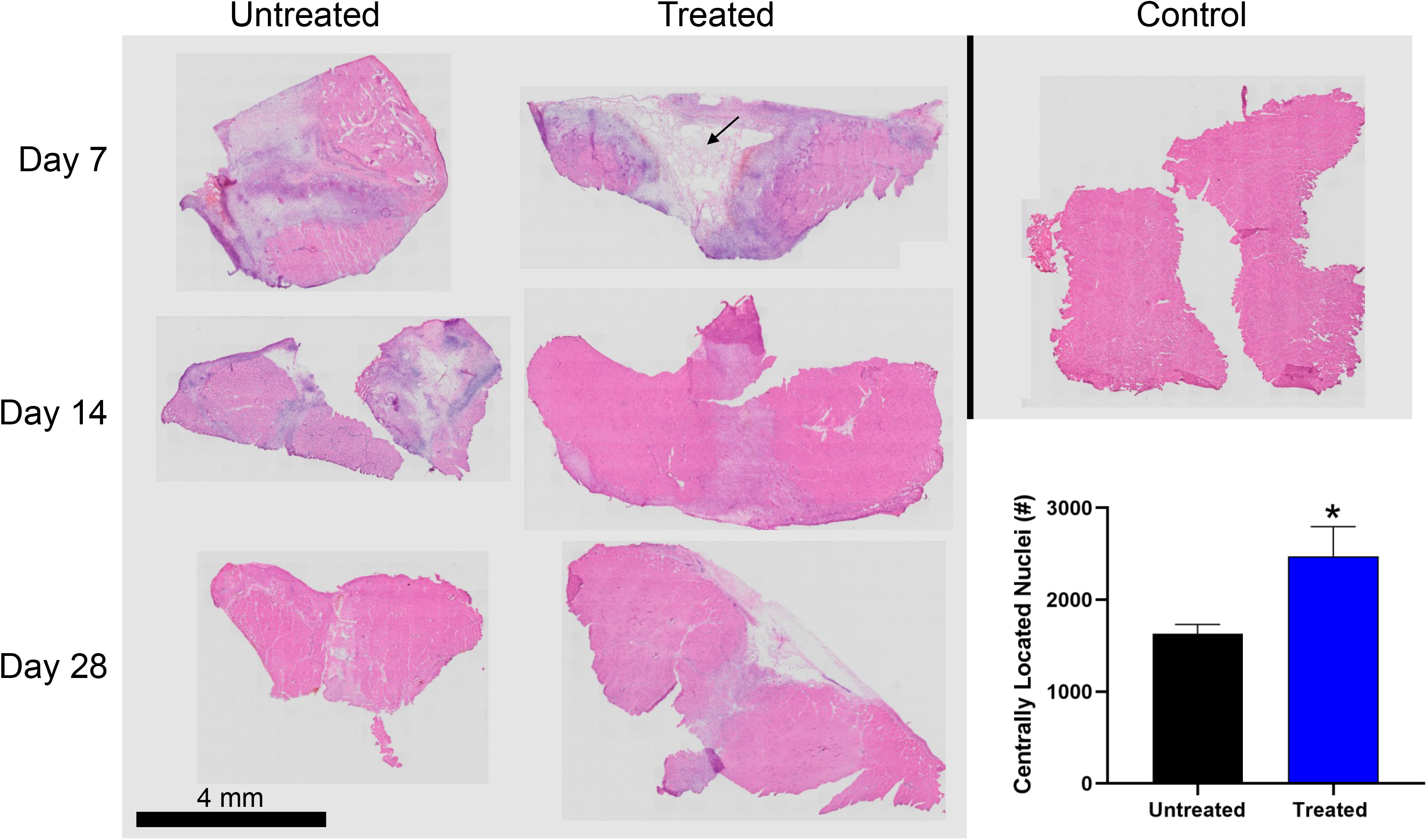
Transverse muscle cross-sections were stained with H&E and whole slide images were captured at days 7, 14, and 28 post-injury. Treated muscles had more myofibers with centrally located nuclei at day 28. “*” indicates a statistical difference (p < 0.05) between different treatment groups. The black arrow indicates the sponge in the defect.

Muscle cross-sections were also stained with myosin heavy chain (MHC) and collagen (COL), as shown in Figure 8. On day 7, both untreated and treated muscles showed a large defect region filled with collagenous fibrotic tissue. The remodeled sponge could be identified in the defect. On day 14, the untreated muscles continued to show collagenous fibrotic tissue deposition. However, in the sponge treated muscles, the defect region appeared smaller with less fibrotic tissue. Several small diameter MHC^+^ myofibers could also be identified in the defect region of the sponge treated muscles (Supplemental Figure 2). By day 28, the sponge treated muscles continued to show a smaller fibrotic region as well as increased presence of MHC^+^ myofibers in and around the defect region. Muscle regeneration and fibrosis was assessed by analyzing the ratio of myosin heavy chain to collagen (MHC: COL). The MHC: COL ratio was statistically similar between the untreated and sponge treated muscles on day 7. On day 14, sponge treated muscles trended towards an increased MHC: COL ratio (p=0.055), and on day 28, the sponge treated muscles showed statistically higher MHC: COL ratio compared to the untreated muscles.

**Figure 8.**
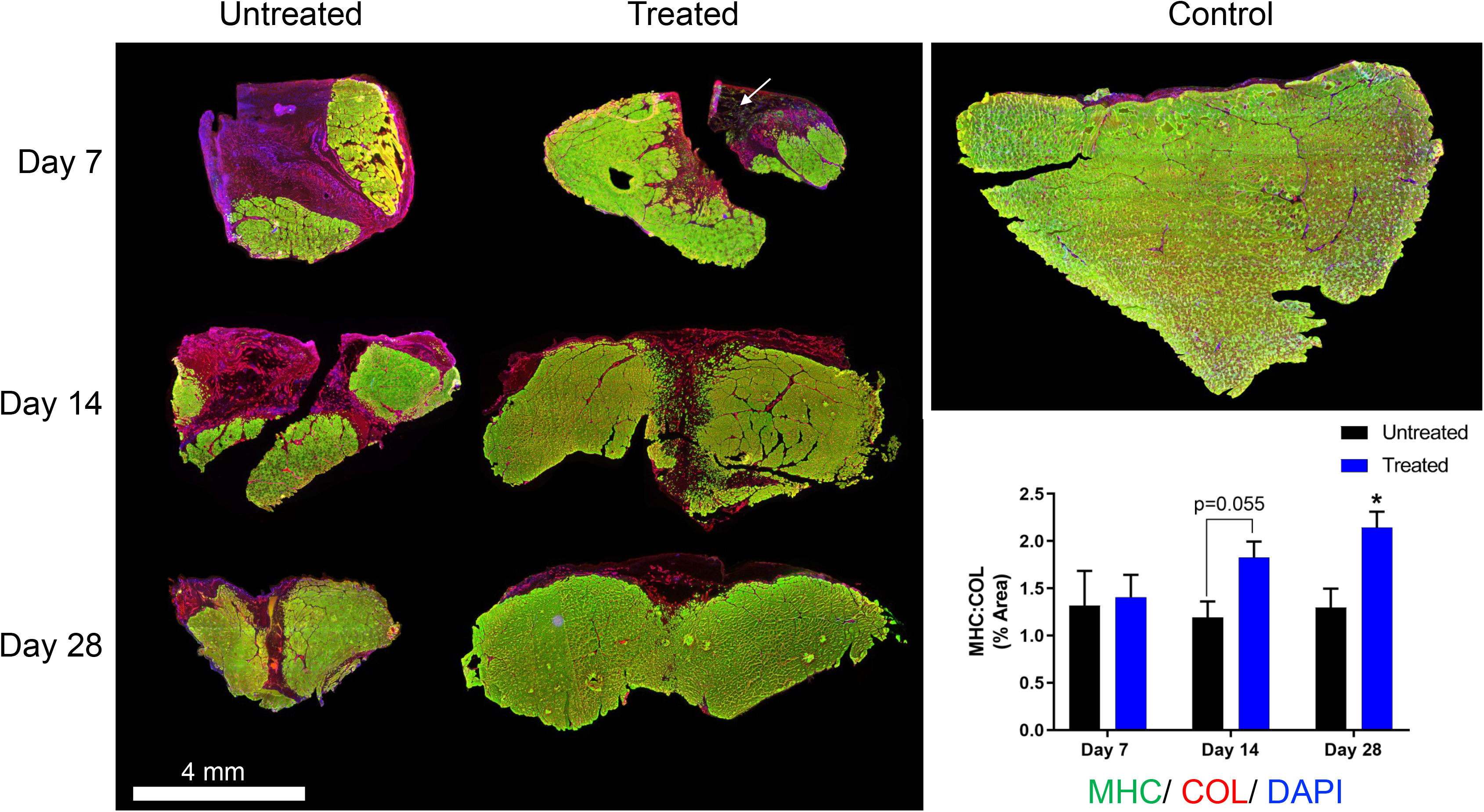
Transverse muscle cross-sections were stained with myosin heavy chain (MHC), collagen (COL), and DAPI. The % area of MHC and COL were quantified and MHC:COL ratios were calculated. Treated muscles have higher MHC:COL ratios by day 28 post-injury. “*” indicates a statistical difference (p < 0.05) between different treatment groups at a particular time-point. The white arrow indicates the sponge in the defect.

Qualitative analysis of the cellular infiltration showed that while CD3^+^ positive cells remained around the periphery, the CD68^+^ macrophages infiltrated the three-dimensional structure of the sponges (Figure 9). Both untreated and treated muscles supported CD31^+^ endothelial cells and SCA1^+^ stem cell activity in the VML defect region.

**Figure 9.**
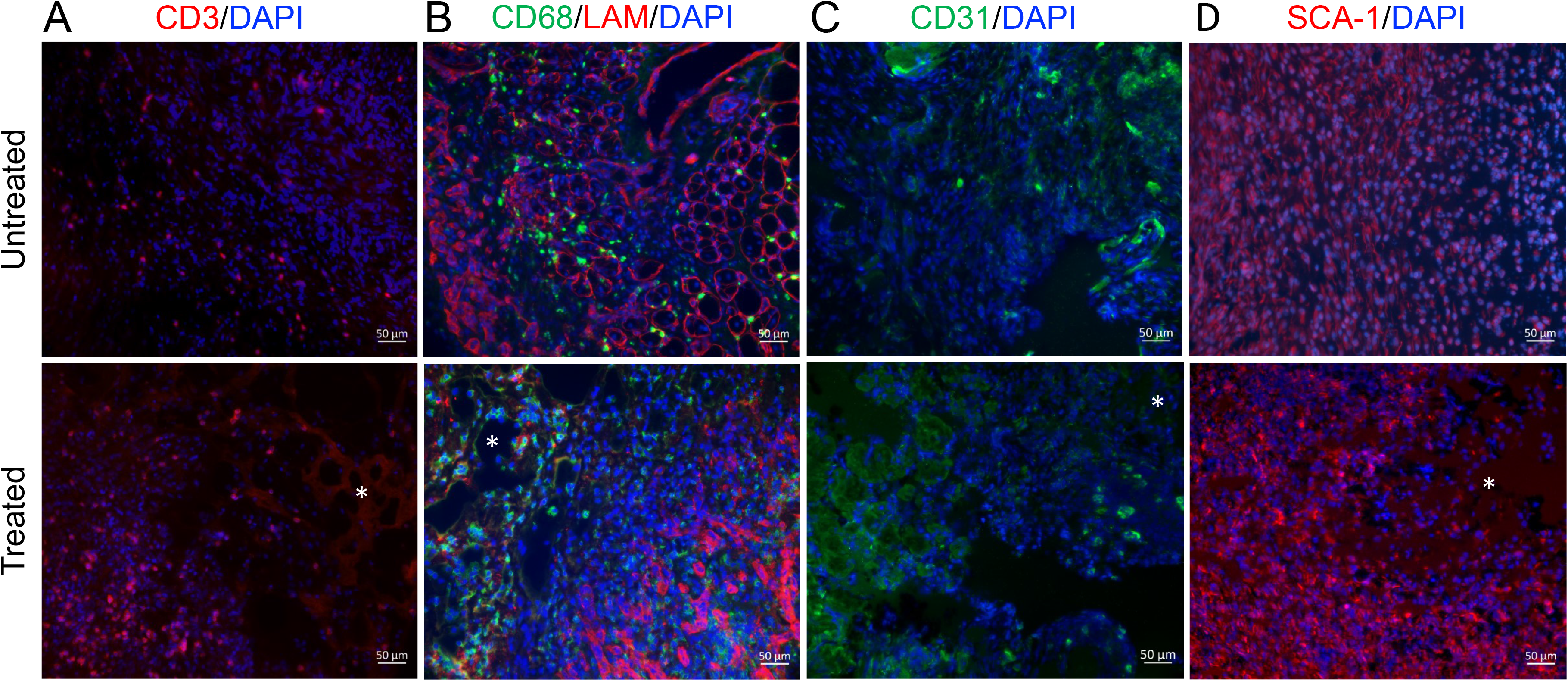
Muscles were stained for (A) T-cells (CD3), (B) macrophages (CD68), (C) endothelial cells (CD31), and (D) stem cells (SCA-1). The white asterisk indicates the sponge in the defect.

### Cross-sectional area measurement

The total number of myofibers in the cage control TA muscle cross-sections were determined to be ~9865. VML injury reduced the total number of myofibers in the TA muscle by 76.5% on day 14. On both days 14 and 28, sponge treatment significantly increased the total number of myofibers (treatment effect, p=0.0205). The percentage improvement in the number of myofibers with sponge treatment was 77% and 82%, on days 14 and 28, respectively. The mean and median CSA was statistically similar between the untreated and treated muscles. The distribution of fiber CSA is shown in Figure 10. Sponge treatment significantly increased the number of small diameter myofibers (<500 μm^2^) on both days 14 and 28 post-VML injury. The number of fibers in the CSA range of 500-999 μm^2^ and 1000-1499 μm^2^ were also significantly higher with sponge treatment on day 14 post-VML. The number of small diameter fibers (<500 μm^2^) showed a significant increase from day 14 to day 28 in the sponge treated groups. A simultaneous decline in the number of bigger fibers (500-1500 μm^2^) was observed with sponge treatment on day 28 compared to day 14.

**Figure 10.**
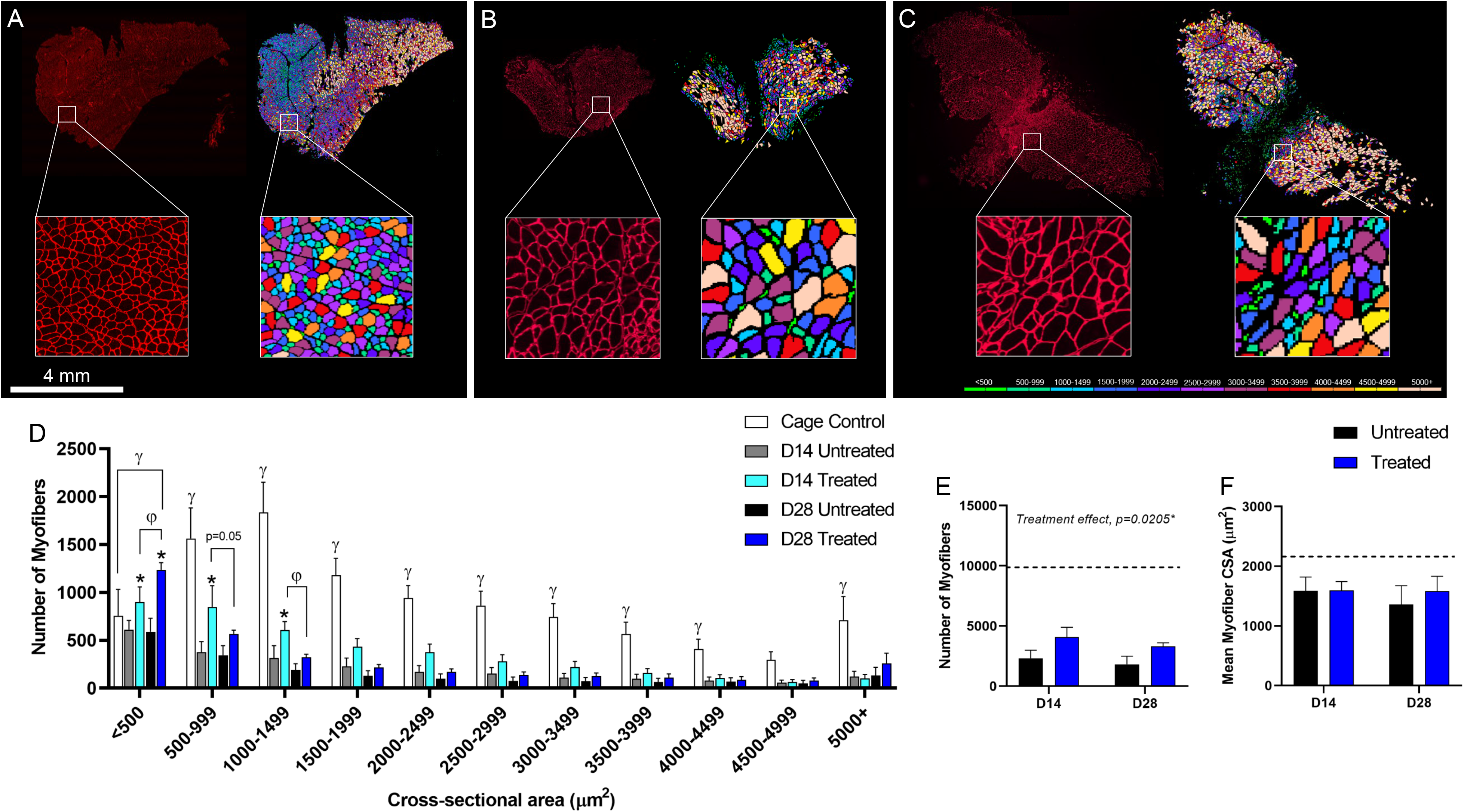
Muscle fibers were outlined with laminin and cross-sectional areas were quantified. Myofiber CSA at day 28 for (A) cage control, (B) untreated, and (C) treated groups are displayed as color-coded maps. (D) A higher quantity of small CSA fibers (<500 μm^2^) were present in treated muscles at both days 14 and 28 post-injury. (E) Treated muscles also contained higher numbers of total myofibers than untreated muscles at days 14 and 28. (F) The mean CSA was not different between untreated and treated groups. “*” indicates a statistical difference (p < 0.05) between untreated and treated muscles at a particular time-point. “γ” indicates a statistical difference (p < 0.05) between controls and injured muscles at a particular time-point. “φ” indicates a statistical difference (p < 0.05) between day 14 and 28 for a particular treatment group. The dashed line indicates control.

Fiber type specific CSA analysis revealed no change in the percentage of either Type 1, 2A, 2B or 2X myofibers with VML injury (Figure 11). However, the mean CSA of fast-twitch glycolytic fibers (i.e., type 2B and 2X) was significantly lower with VML injury. Sponge treated muscles showed significantly larger CSA of type 2B myofibers compared to untreated muscles. Fiber type distribution analysis showed that sponge treatment significantly increased the number of small diameter fibers (<500 μm^2^) in the slow oxidative fibers (i.e., type 1 and 2A), and large-diameter myofibers (>4000 μm^2^) in the fast-twitch type 2B fibers.

**Figure 11.**
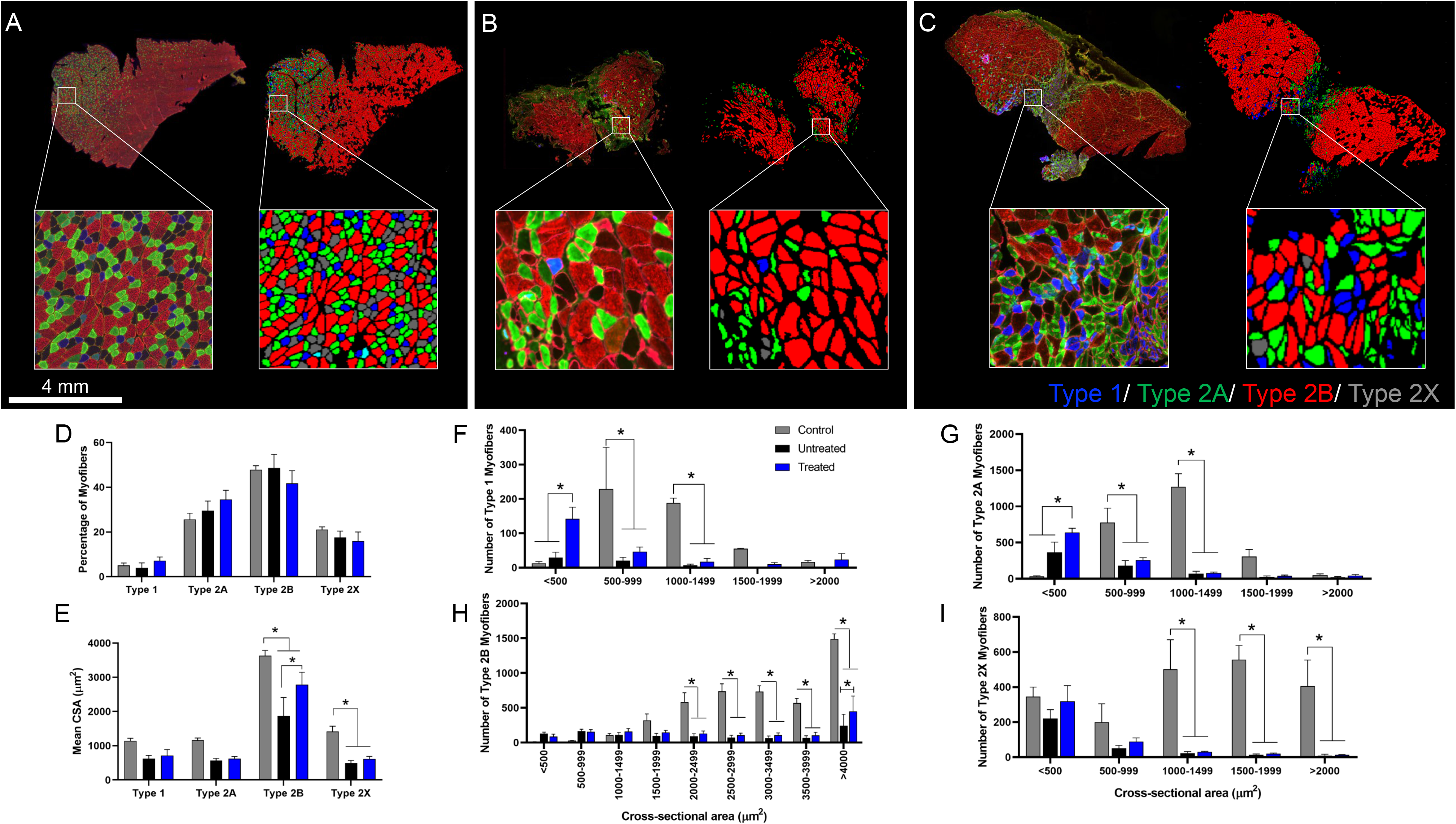
Muscle cross-sections were stained for different fast- and slow- twitch fiber types. Immunostained images and color-coded maps are displayed for (A) uninjured controls, (B) untreated muscles, and (C) sponge-treated muscles. Fiber type distribution and CSA was quantified. (D) VML had no effect on the percentages of different types of fibers, but (E) decreased the mean CSA of fast-twitch fibers (i.e., type 2B and 2X). Sponge treatment increased the mean CSA of Type 2B fibers compared to untreated muscles. Fiber type CSA distributions for (F) Type 1, (G) Type 2A, (H) Type 2B, and (I) Type 2X myofibers are shown. “*” indicates a statistical difference (p < 0.05) between different treatment groups.

## Discussion

The most important finding of this study is that FK-506 loaded biomimetic sponges improve muscle structure and function in a rodent model of VML. Sponge treatment increased the total number of myofibers, type 2B fiber CSA, MHC: collagen ratio, and muscle function compared to untreated VML injured muscles. These results are encouraging, and we believe that biomimetic sponges may provide a promising “off-the-shelf” approach for VML that is worthy of clinical investigation.

An increase in the number of myofibers and type 2B fiber CSA may underlie the basis for improved muscle function with sponge treatment. To the best of our knowledge, a significant increase in myofiber number and muscle function has never been reported with the implantation of an acellular ECM scaffold in a full-thickness VML model. The use of decellularized scaffolds in VML injuries have shown conflicting results with studies reporting either no improvement [15, 29, 30], exacerbated force deficits [31], or modest improvements in force production [32]. While scaffolds that contain myogenic stem cells have shown improvement in muscle regeneration and force production [5, 33–35], their clinical translation is likely to be hindered by limited availability of autologous donor tissue and potential donor-site morbidity.

In this study, the biomimetic sponges were expected to provide both immunomodulatory as well as regenerative effects. The sponges contain an optimized ratio of ECM proteins such as gelatin, collagen, and laminin-111, and have supported myogenic activity *in vitro* and *in vivo* in a previous study [25]. Besides FK-506, immunomodulatory and anti-inflammatory effects can also be ascribed to ECM scaffolds [36]. While it is difficult to attribute specific immunomodulatory or regenerative effects to the different constituents of biomimetic sponges, our results suggest that their implantation modulated the VML microenvironment in a way that improved muscle structure and function. In support of this contention, the flow cytometry data showed that the implantation of sponges changes the temporal dynamics of the immune response in VML injured muscles. In contrast to untreated muscles where the presence of T-cells (CD3^+^, CD4^+^) significantly reduces between days 7-14, the sponge treated muscles show maintained T-cell presence over 14 days post-injury. Interestingly, the percentage of CD8^+^ T-cells peaks at day 14 in sponge treated muscles while it remains unchanged between days 7-28 in untreated muscles. At day 7 post-injury, while the percentage of infiltrating CD8^+^ cells was unaffected, the gene expression of CD8a was significantly lower in sponge treated muscles.

To gain further insight into the immune response within the first 7 days post-injury, we performed a PCR immunoarray. Of the 87 genes related to innate and adaptive immune responses, 35 genes in untreated group and 28 genes in treated group were significantly higher than the control group. The innate immune response is the first line of defense and is initiated by damage associated molecular patterns (DAMPs) such as cellular fragments or degraded ECM proteins that can stimulate TLRs and NLRs. The gene expression of TLRs-1, 5, 6 and 9 was significantly higher with sponge treatment compared to uninjured controls. TLRs have been implicated in the recognition of implanted biomaterials [27]. Besides antigen presenting cells of the innate immune system, the expression of TLRs has been reported on regenerating myofibers and vascular endothelial cells in inflammatory myopathies [35]. However, the localization of TLRs was not investigated in this study.

The complement system is a crucial component of innate immunity [37]. In this study, the gene expression complement opsonin C3 was significantly higher in the sponge treated muscles compared to uninjured controls. It has been shown that implantation of biomaterials can activate the complement system [38]. Besides initiating inflammation, the complement system also plays a role in muscle repair. For instance, muscle regeneration was found to be impaired following cardiotoxin injury in C3-deficient mice due to lower macrophage infiltration and decreased myoblast proliferation [39].

The implantation of FK-506 loaded sponges resulted in the downregulation of several inflammatory genes associated with T-cell activity such as CD8a, CXCL10, CXCR3, CD40, caspase 1, and FasL. Studies have shown that FK-506 can inhibit CXCL10 [40], CD8+ T-cell expansion, and FasL activity [41]. T-cells express CXCR3, a receptor for CXCL10. Binding of CXCL10 to CXCR3 initiates inflammation, promotes apoptosis, and reduces angiogenesis [40, 42–44]. CXCL10-CXCR3 axis is also important for peripheral homing of T-cells, and the gene expression of both these molecules was downregulated with sponge treatment on day 7. T-cell activation requires CD40 ligand (CD40Lg) binding to CD40, which is expressed on antigen-presenting cells. It has been suggested that CD40-CD40Lg binding can directly and independently co-stimulate T cells [45]. The blockade of this interaction resulted in long-term cardiac allograft survival in a murine model, suggesting that inhibiting CD40 alone can suppress T-cell mediated immunogenic responses. The cytotoxic activity of CD8 cells is mediated by apoptosis-associated genes such as caspase 1 and FasL. Interestingly, FasL has been implicated in muscle atrophy and degeneration in Duchenne muscular dystrophy (DMD) patients [46].

Sponge treatment also resulted in significant upregulation of interferon-γ (IFN-γ). IFN-γ is primarily secreted by activated T-cells. It is an inflammatory cytokine that regulates various immune responses and physiological processes [47–49]. Inflammatory myopathies typically show increased type 1 interferon signaling and elevated expression of type I IFN stimulated genes (ISG) such as MX1, CXCL10, and IRF7 [50]. Atrophic small diameter myofibers in dermatomyositis patients have shown co-expression of atrogin 1 and MX1. A persistent IFN-induced response was implicated in the pathophysiology of the juvenile dermatomyositis, which is characterized by necrotic pathways and degeneration/regeneration cascades [51]. In sponge treated VML injured muscles, IFN-γ was upregulated while ISGs such as MX2, CXCL10, and IRF7 were downregulated. The receptor, IFN-γR1, was also found downregulated in response to sponge treatment. These results suggest a potential negative regulation of IFN-γ signaling by sponge treatment [52]. Several desensitization pathways for IFN-γ signaling have been proposed [53]. However, more studies are needed to determine if ISGs exacerbate VML injury similar to inflammatory myopathies. While FK-506 has been effective against inflammatory myopathies in several studies [54, 55], it is unclear if it inhibits ISGs or IFN-γ signaling in muscle injuries such as VML.

Taken together, FK-506 loaded sponges have an overall negative impact on T-cell associated gene expression within the first week post-injury, but this effect is short-lived. The peak in T-cell quantity on day 14 post-injury was almost synchronous with the clearance of FK-506. Coinciding with an increased T-cell presence on day 14, an increase in the number of small diameter (<500 μm^2^) myofibers, and MHC: COL ratio was also observed. T-cells play complex roles in skeletal muscle regeneration and show varied responses to muscular dystrophies and injuries. Pathological muscle conditions such as Duchenne Muscular Dystrophy or polymyositis are characterized by the increased and persistent presence of CD8^+^ T −cells [25]. Studies have suggested that CD8+ T cells could have a direct cytotoxic role on muscle fibers expressing major histocompatibility complex (MHC) class I molecules [56]. Healthy individuals do not show MHC class I myofibers, but they are frequently observed in myositis patients. In another study, depletion of CD8^+^ T-cells in cardiotoxin damaged Casitas B-lineage lymphoma-b (Cbl-b)-deficient mice resulted in improved regenerative outcomes [26].

In contrast, other studies have indicated that T-cells might play a role in muscle regeneration, and completely abolishing T-cell response could have detrimental effects. For instance, in a model of cardiotoxin induced muscle injury, it was observed that in the absence of CD8^+^ T-cells, matrix deposition is increased. At the same time, monocyte recruitment, myoblast proliferation, and myofiber growth are diminished [57]. In a study by Hurtgen *et al.*, implantation of minced muscle autografts resulted in enhanced presence of CD3^+^, CD4^+^, and CD8^+^ T cells in a VML model [58]. The increased presence of these cell types over 14 days post-injury did not hinder muscle regeneration, as evidenced by newly regenerating myofibers and improved functional recovery. In a human study, increased presence of CD8^+^ T-cells was implicated in muscle adaptation to repeated eccentric contractions [59, 60]. The fact that CD8^+^ T-cells were increased significantly only after second bout of exercise when evidence of muscle damage was reduced, suggested that these cells do not exacerbate injury but facilitate repair [59, 61]. Therefore, it appears that increased CD8^+^ T-cell infiltration coincides with muscle repair, but the exact mechanism through which CD8^+^ T-cells participate in muscle regeneration remains unknown and needs to be investigated in future studies.

Interestingly, we did not observe heightened myogenic gene or protein expression with sponge treatment of VML injury. However, quantitative analysis showed a significantly higher number of small diameter myofibers (<500 μm^2^) with treatment. We believe that myofiber splitting could account for the significant increase in the small diameter myofibers on days 14 and 28 in the sponge treated muscles [62]. The leftward shift in the muscle fiber size distribution curve (Supplementary Figure 3) between days 14 and 28 suggests that larger myofibers may have split during this period. Qualitative analysis showed several myofibers with irregular shapes and displaced myonuclei, from which a smaller myofiber appeared to have broken apart (Supplementary Figure 3). Myofiber splitting has been observed in hypertrophy models [63], which indicates that the process may be an adaptation to maintain either the myonuclear domain, oxygen diffusion capacity, or force production [64]. In VML models, the remaining muscle mass experiences chronic overload as it attempts to compensate for the lost tissue [8]. It has been suggested that myofiber splitting in response to increased loading can be biomechanically advantageous as it distributes the force over a larger surface area [64]. Therefore, myofiber splitting may account for higher myofiber counts and peak isometric torque in sponge treated muscles.

In this study, the sponge treated muscles showed several clusters of small-diameter myofibers, the majority of which appeared to be Type 1 or 2A myofibers. Quantitative analysis also confirmed significantly higher numbers of small diameter myofibers (<500 μm^2^) in the fiber type 1 and 2A category. Taken together, these results might indicate that the slow oxidative fiber types are more prone to splitting in the VML model. We have also demonstrated that VML injury primarily impacts the CSA of fast glycolytic fibers (i.e., Type 2B and 2X). This result is in agreement with previous studies where eccentric contractions [65] and DMD [66] were shown to damage type 2 fibers selectively. Sponge treatment partially rescued the CSA of type 2B but not 2X myofibers. Sponge treated muscles showed significantly higher CSA of type 2B myofibers as well as greater number of large diameter (> 4000 μm^2^) type 2B fibers compared to untreated VML injured muscles. An improvement in type 2B myofiber CSA could also account for increased force production. These findings have major implications for VML injured patients because large type 2B fibers can withstand substantial loads. Therefore, preventing their loss and preserving their CSA might help improve the quality of life in affected patients [67].

Overall, these results suggest that implantation of biomimetic sponges protects the remaining muscle mass from necrosis and degeneration. Sponge treatment appears to reduce the severity of the injury and allows for improvements in muscle structure and function. These results could be attributed to FK-506 driven immunomodulation, replenishment of vital ECM proteins, and mechanical support offered by the three-dimensional scaffold. A potential limitation of this study is the rapid release of the unconjugated FK-506 that was encapsulated in the biomimetic sponges. FK-506 showed a rapid burst release *in vitro* as it diffused out of the sponges. We did not investigate the *in vivo* release kinetics of FK-506, but it can be reasonably assumed that the drug is cleared within the first few days after sponge implantation. Interestingly, it has been shown that dendritic cells can sequester FK-506 and release it slowly in quantities that can inhibit T-cell activity for at least 72 hours [68]. Dendritic cells have been detected as early as 24 hours after muscle injury [69, 70]. While continuous administration of FK-506 has been done in VML models [23, 24], others have reported that a single local application of FK-506 at the time of repair can support functional nerve regeneration 2-3 months after injury [71, 72]. Future studies may investigate the impact of sustained-release FK-506 on VML injury to determine if a short-term administration is adequate. However, in a clinical setting, prolonged immunomodulation can be easily achieved by administering FK-506 either as an oral dose or intravenous infusion [73]. Future work will also investigate the extent to which functional recovery can be enhanced by combining biomimetic sponge treatment with physical rehabilitation [74].

## Figure Captions

**Supplemental Figure 1.**
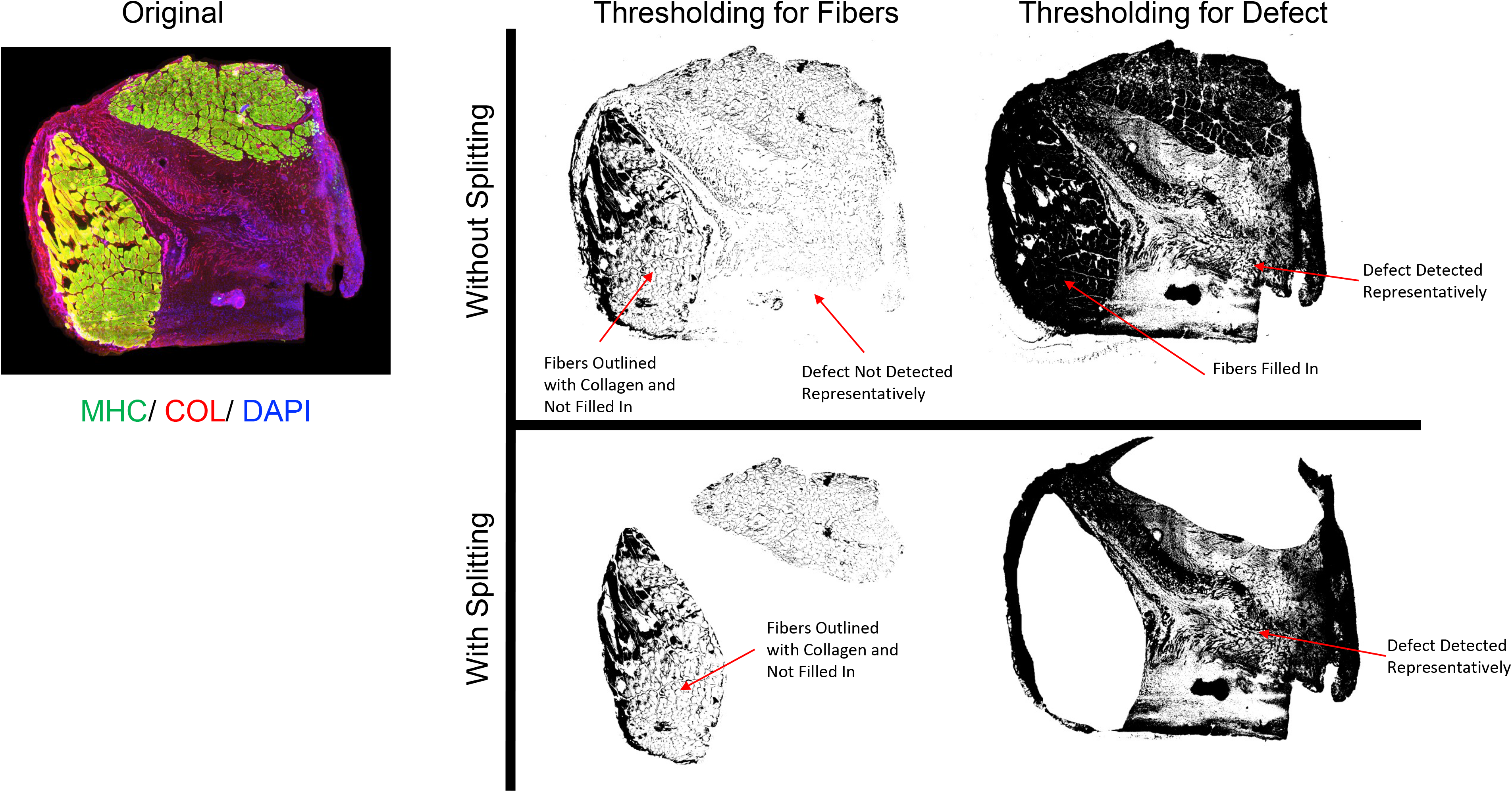
Full-scale images of muscle sections stained with MHC and COL were used to quantify the percent area of each. Thresholding without separating the remaining muscle tissue and the defect region (top) results in a less representative collagen % area. By contrast, splitting the remaining muscle tissue and defect region into two separate images (bottom) results in higher accuracy because the defect can be detected without muscle fibers being filled in with collagen.

**Supplemental Figure 2.**
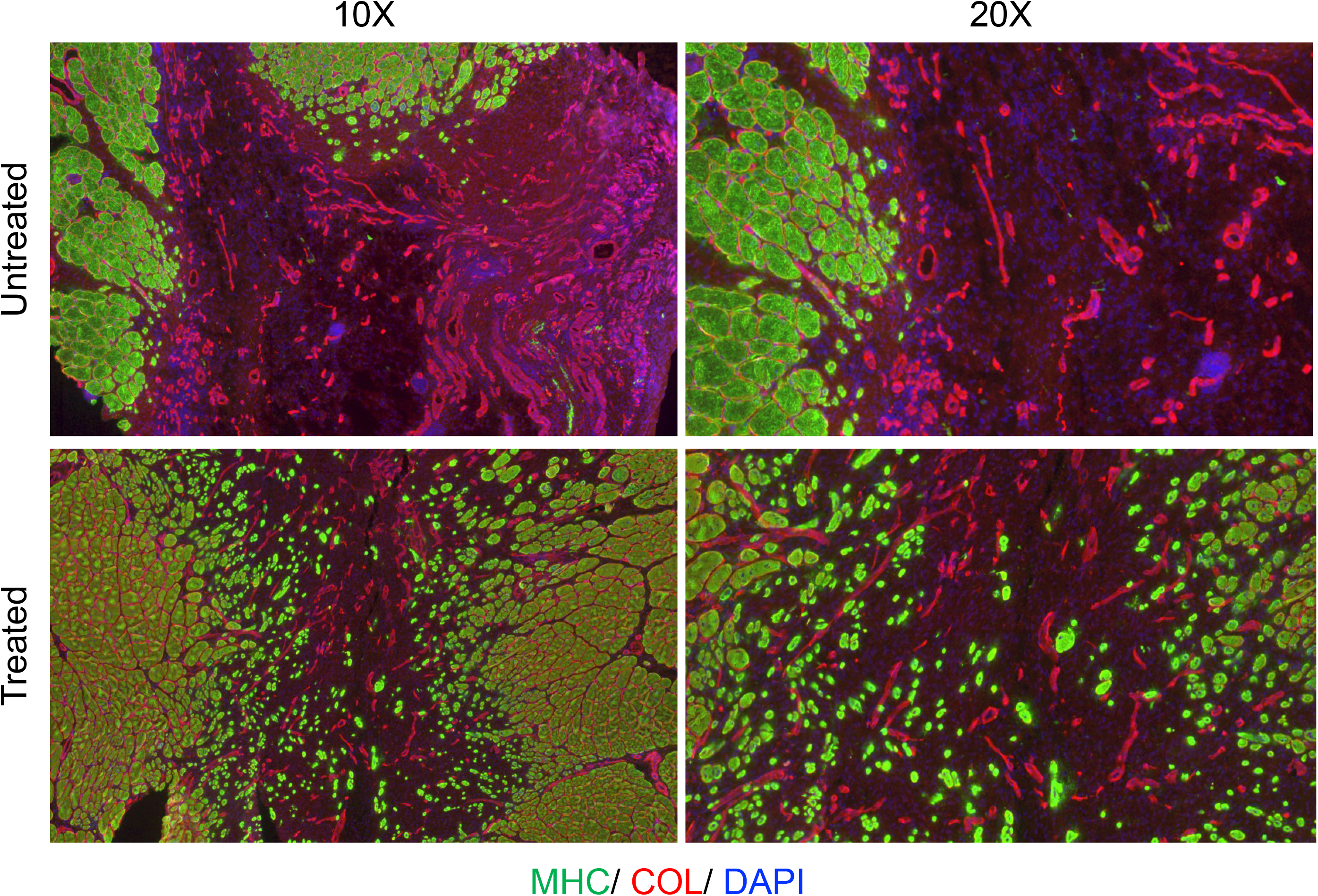
Higher magnification images provide a closer look at the defect region. Sponge treated muscles show more small-diameter myosin^+^ fibers at day 14 post-injury.

**Supplemental Figure 3.**
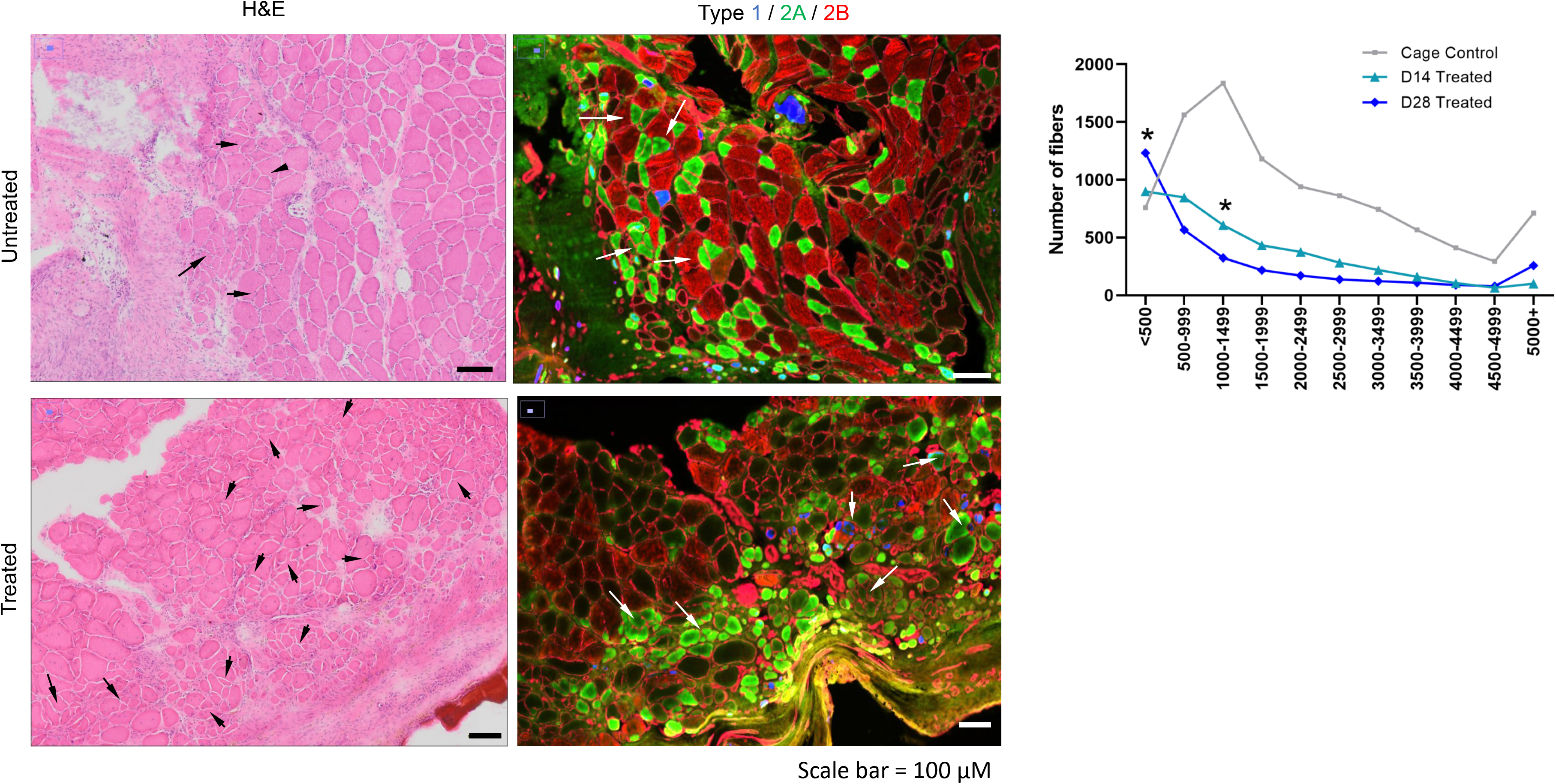
Qualitative analysis of split myofibers is presented. Arrows show the branched/split myofibers. Muscle cross-sectional areas are represented as a fiber size distribution curve. “*” indicates a statistical difference (p < 0.05) between different groups.

## Acknowledgments

This work was supported by a grant from the National Institute of Health (NIGMS) 1R15GM129731 awarded to KG. We would like to thank Joy Eslick and Sherri Koehm for help with the flow cytometry experiments. We would like to thank Gary D. London (Washington University), as well as Caroline Murphy and Dr. Grant Kolar (Saint Louis University) for technical assistance with histological imaging.

## Conflict of Interest

KG has equity interest in GenAssist, Inc., and serves on the company’s scientific advisory board. GenAssist, Inc. is developing products related to the research described in this paper. The terms of this arrangement have been reviewed and approved by Saint Louis University, in accordance with its conflict of interest policies. GJH is the CEO and JM is the CTO of GenAssist, Inc. and both are members of the board of directors. AJD also holds equity interest in GenAssist, Inc.

## Notes

### Competing Interest Statement

The authors have declared no competing interest.

